# The impact of size on particle drainage dynamics and antibody response

**DOI:** 10.1101/2020.09.28.316612

**Authors:** Simon Zinkhan, Anete Ogrina, Ina Balke, Gunta Reseviča, Andris Zeltins, Simone de Brot, Cyrill Lipp, Xinyue Chang, Lisha Zha, Monique Vogel, Martin F. Bachmann, Mona O. Mohsen

## Abstract

Vaccine-induced immune response can be greatly enhanced by mimicking pathogen properties. The size and the repetitive geometric shape of virus-like particles (VLPs) influence their immunogenicity by facilitating drainage to secondary lymphoid organs and enhancing interaction with and activation of B-cells and other innate humoral immune components. VLPs derived from the plant Bromovirus genus, specifically cowpea chlorotic mottle virus (CCMV), are T=3 icosahedron particles. They can be easily expressed in an *E*. coli host system and package ssRNA during the expression process. Recently, we have engineered CCMV-VLPs by incorporating the universal tetanus toxoid (TT) epitope at the N-terminus. The modified CCMV_TT_-VLPs successfully form icosahedral particles *T=3*, with a diameter of ∼30nm analogous to the parental VLPs. Interestingly, incorporating TT epitope at the C-terminus of CCMV_TT_-VLPs results in the formation of Rod-shaped VLPs, ∼1µm in length and ∼30nm in width. In this study, we have investigated the draining kinetics and immunogenicity of both engineered forms (termed as Round-shaped CCMV_TT_-VLPs and Rod-shaped CCMV_TT_-VLPs) as potential B cell immunogens using different *in vitro* and *in vivo* assays. Our results reveal that Round-shaped CCMV_TT_-VLPs are more efficient in draining to secondary lymphoid organs to charge antigen-presenting cells as well as B-cells. Furthermore, compared to Rod-shaped CCMV_TT_-VLPs, Round-shaped CCMV_TT_-VLPs led to more than 100-fold increased systemic IgG and IgA responses accompanied by prominent formation of splenic germinal centers. Round-shaped CCMV_TT_-VLPs could also polarize the induced immune response towards TH_1_. Up to our knowledge, this is the first study investigating and comparing the draining kinetics and immunogenicity of one and the same VLP monomer forming nano-sized icosahedrons or rods in the micrometer size.

## Introduction

In 1956, Crick and Watson have stated that “it is a striking fact that almost all small viruses are rods or spheres”, “These shells are constructed from a large number of identical protein molecules, of small or moderate size, packed together in a regular manner” (1). The main reason for this arrangement is the small genome of viruses, especially RNA viruses. The coat protein (CP) of many viruses is made up of multiple copies arranged in an icosahedron or a helical-shaped geometry (2, 3). The icosahedral structure of viruses is more prevalent than the helical-shaped one.

Virus-like particles (VLPs) have emerged in the last few decades as a premium vaccine platform due to several reasons including: being a safe platform lacking replicating genetic materials, their repetitive surface geometry that serves as a potent pathogen-associated structural pattern (PASP), their ability to package different toll-like receptor ligands (TLRs), the feasibility in coupling different epitopes and most importantly their favorable size ranging between 20-200nm (4). Such size allows VLPs to rapidly and efficiently filter and drain through the conduit system and gain access to lymphoid follicles (4-7). The approved VLP-based vaccines currently on the market mostly exhibit an icosahedral surface geometry based on the quasi-equivalence concept described by Caspar and Klug in 1962 and expressed as *Triangulation (T)* (8, 9). For example, human papilloma viruses (HPVs) are *T=7* of ∼60nm in size (10), while hepatitis E virus (HEV) VLPs are *T=1* of ∼25nm (11). The different generations of hepatitis B virus (HBV) vaccines show highly organized sub-viral particles (SVPs) of ∼20-25nm (12). In contrast, the arrangement of CPs of VLPs in helical or rod-shape geometry is also possible; tobacco-mosaic virus (TMV) is a well characterized representative of this category. Virions of TMV are ∼300nm in length and ∼18nm in width (13, 14). TMV-VLPs have been investigated as a promising platform in nanotechnology (15) and as a vaccine platform as well (16). Nevertheless, knowledge is scarce regarding TMV-VLPs drainage dynamics. Icosahedral VLPs can be manipulated by inserting few mutations to form rod-shaped VLP. For example, VLPs derived from the bacteriophage Qβ can assemble in a rod-shaped particle following the mutation of five amino acid (a.a.) residues in the FG loop of its CP (17).

It is known that the repetitive surface geometry of icosahedral VLPs enhances optimal induction of B-cell response via cross-linking of B-cell receptors (BCRs) (18, 19). Previously, we have shown that displaying epitopes on icosahedral *T=3* VLPs such as bacteriophage Qβ or plant-derived CuMV_TT_ VLPs result in high specific IgG antibody (Ab) titers as well as neutralizing Abs (20-25). Some studies have revealed that a vaccine based on rod-shaped tobacco-mosaic (TMV)-VLPs could also serve as an effective platform to display different epitopes capable of eliciting an immune response against different pathogens (26, 27).

Cowpea chlorotic mottle virus (CCMV) is a Bromovirus naturally infecting plants and therefore is non-infectious to humans. The infected plants develop yellow spots on their leaves, hence termed chlorotic (28). The virus is an icosahedron *T=3* of ∼28nm in diameter, consisting of 180 sequentially identical CPs. The coat protein can adopt multiple quasi-equivalent forms referred to as the coat subunits A, B and C forming either hexamers (alternating subunits B and C) or pentamers (subunit A). The resulting virus particle consists of 12 pentamers and 20 hexamers (29). Previously, it has been shown that icosahedral CCMV can be converted into rod-shaped particles after a disassembly/reassembly process (30).

In this study, we have demonstrated that CCMV-derived VLPs of different morphology (icosahedral or rod-shaped structure) can be obtained directly from recombinant *E*.coli cells. We specifically manipulated CCMV-VLPs by inserting the universal tetanus toxoid (TT) epitope at the C or N-terminus to form icosahedral VLPs in the nanometer scale or rod-shaped VLPs with sizes in the micron range. Such intervention allowed us to study the impact of size on particles in terms of drainage dynamic and magnitude of induced immune response using VLPs based on essentially the same VLP-monomer. Our results demonstrate for the first time that VLPs in the nm size range are vastly more immunogenic than micron-sized particles.

## Materials and methods

### Round-shaped and Rod-shaped CCMV_TT_-VLPs cloning, expression and production

Cloning of CCMV-CP with induced tetanus toxoid epitope for expression: A cloned copy of the CCMV coat protein gene (wt *CP)* was obtained from Dr. Alain Tissot (Zürich) and used in PCR mutagenesis for insertion of the coding sequence of tetanus toxoid epitope (TT830 – 843; QYIKANSFIGITE) in 5’- and 3’-terminal ends of the CP gene. To replace the original amino acids at the N-terminus of CCMV CP with the TT epitope sequence, the pET42-CCMVwt plasmid was used as a template for PCR amplification and mutagenesis. *NdeI* site at the 5’end of the CCMVwt gene was used for cloning corresponding PCR products. To introduce the tetanus toxoid epitope coding sequence into the CCMV-wt gene, two step PCR mutagenesis was necessary. For the first step amplification the following primers were used for N terminal PCR: CC_N83_d24F (5’ATACATATGGGCCAGTATATTAAGGCCAACTCCAAATTTATCGGGATTACCGA 3’) and CC_N83R (5’ AGTTAACTTCCCTGTACCGACTGTTTCGGTAATCCCGATAAATTTGGAGTTG 3’). For the second round the PCR products from the first round were diluted 1:50 and re-amplified with primers CC_N83_d24F and CC_AgeR (5’ ACTTCGATACGCTGTAACCGGTCCA 3’). For C terminal insertion of TT epitope the following primers were used: CC_C83F (5’ TGACGACTCTTTCACTCCGGTCTATGGCCAGTATATTAAGGCCAACTCC 3’) and CC_C83R (5’ TTACTCGAGAAGCTTATTACTCGGTAATCCCGATAAATTTGGAGTTG 3’). Second round of the PCR was performed as describe above using primers CC_C83R (5’ TTACTCGAGAAGCTTATTACTCGGTAATCCCGATAAATTTGGAGTTG 3’) and CC_SacIIF (5’ CCCTTGAACAACTCGCCGCGGA 3’).

The corresponding PCR fragments were analysed in 0.8% agarose gel and then purified using the *GeneJet Gel Extraction* kit (Thermo Fischer Scientific, Waltham, Massachusetts). Then the 5’ terminal end PCR product and plasmid pET42-CCMVwt were digested with enzymes *NdeI* and *BshTI* (Thermo Fischer Scientific, Waltham, Massachusetts) and ligated, resulting in plasmid pET42-CCMV-Ntt830. The 3’terminal end PCR product and plasmid pET42-CCMVt were digested with enzymes *Cfr42I* and *XhoI* (Thermo Fischer Scientific, Waltham, Massachusetts) and ligated, resulting in plasmid pET42-CCMV-Ctt830.

*E.coli* XL1-Blue cells were used as a host for cloning and plasmid amplification. To avoid PCR errors several CP gene-containing pET42 plasmid clones were sequenced using the BigDye cycle sequencing kit and an ABI Prism 3100 Genetic analyzer (Applied Biosystems, Carlsbad, USA). After sequencing, the plasmid clones without sequence errors were chosen for further experiments.

Cloning of CCMV-SS-CP with induced tetanus toxoid epitope for expression: To obtain “salt-stable” CCMV VLPs, the replacement of lysine against arginine in position 42 (K42R) was necessary. For CCMVwt-SS the following primers were used: CCP_salt_AgeI_R (5’ TGTAACCGGTCCATGCTTTAATAGCGCGGCCTT 3’) and CCM_NdeF (5’ ATACATATGTCTACAGTCGGTACAGGG 3’). For CCMV-Ntt830-SS the following primers were used: CC_N83_d24F and CC_salt_AgeI_R. The corresponding PCR products were cloned into the pTZ57R/T vector (Thermo Fischer Scientific, Waltham, Massachusetts). *E. coli* XL1-Blue cells were used as a host for cloning and plasmid amplification. To avoid RT-PCR errors, several CP gene-containing pTZ57 plasmid clones were sequenced using the BigDye cycle sequencing kit and an ABI Prism 3100 Genetic analyser (Applied Biosystems, Foster City, California). After sequencing, corresponding DNA fragments without sequence errors were subcloned into the *NdeI/AgeI* sites of pET42-CCMVwt and pET42-CCMV-Ntt830 expression vectors, resulting in the expression plasmids pET42-CCMV-SS and pET42-CCMV-Ntt830-SS. For C terminal tetanus toxoid CCMV-SS construct the corresponding *NdeI/BsrGI*-fragment from pET42-CCMV-SS was subcloned into pET-CCMV-Ctt830, generating the expression vector pET42-CCMV-Ctt830-SS.

Expression and purification of CCMV-SS VLPs: To obtain all salt stable CCMV CP VLPs, each construct was transformed and expressed in *E. coli* C2566 cells (New England Biolabs, Ipswich, USA). After selection of clones with the highest expression levels of the target protein, *E. coli* cultures were grown in 2xTY medium containing kanamycin (25 mg/l) on a rotary shaker (200 rev/min; Infors, Bottmingen, Switzerland) at 30°C to an OD600 of 0.8– 1.0. Then, expression was induced with 0.2 mM Isopropyl-β-D-thiogalactopyranoside (IPTG), and the medium was supplemented with 5 mM MgCl_2_ and 2 mM CaCl_2_. Incubation was continued on the rotary shaker at 20°C for 18 h. The resulting biomass was collected by low-speed centrifugation and was frozen at -70°C. After thawing on ice, the cells were suspended in buffer containing 15 mM sodium phosphate pH 7.5 supplemented with 150 mM NaCl (buffer A) with additional 0.5 mM urea, 1 mM PMSF, 5 mM mercapto-ethanol, and were disrupted by ultrasonic treatment. Insoluble proteins and cell debris were removed by centrifugation (13,000 rpm, 30 min at 5°C). All steps involved in the expression of VLP were monitored by SDS-PAGE using 12.5% gels.

CCMV-SS and CCMV-Ctt830-SS VLPs were separated from cellular proteins by ultracentrifugation (SW28 rotor, Beckman, Palo Alto, USA; at 25,000 rpm, 6 h, 5°C) in a sucrose gradient (20–60% sucrose in buffer A, without mercapto-ethanol and urea, supplemented with 0.5% Triton X-100). The content of gradient tubes was divided into six fractions, starting at the bottom of the gradient, and the fractions were analysed by SDS-PAGE. Fractions containing CCMV-SS CP proteins were combined and dialyzed against 100 volumes of buffer A to remove the sucrose. If necessary, samples were concentrated using an Amicon Ultra-15 centrifugal device (Millipore, Cork, Ireland).

However, soluble proteins of CCMV-Ntt830-SS were precipitated using a mixture of PEG 8,000 (8%) and NaCl (0.15 M), collected by centrifugation and dissolved in buffer A. PEG/NaCl precipitation was repeated for CCMV-Ntt830-SS. After solubilisation or dialysis (in case of CCMV-SS), all CCMV CP preparations were purified two times using an ultracentrifuge and 30% sucrose cushion – first with additional 0.5% Triton X-100 and the second time without Triton X-100 (4 h, 50 000 rmp, 4°C; Type 70Ti rotor, Beckman) and the pellet was then dissolved in buffer A. If necessary, samples were concentrated using an Amicon Ultra-15 centrifugal device (Millipore, Cork, Ireland). To obtain pure preparations of CCMV-CPs for subsequent electron microscopy (EM) analysis, stability and immunological studies, the sucrose gradient, dialysis, and concentration steps were repeated.

All steps involved in the expression and purification of VLP were monitored by SDS-PAGE (using 12.5% gels). The concentration of purified CCMV–CPs were estimated using the QuBit fluorometer in accordance with manufacturer’s recommendations (Invitrogen, Carlsbad, California). Concentrated VLP solutions were stored at +4°C.

### Electron microscopy

Purified Round-shaped or Rod-shaped CCMV_TT_-VLPs proteins (1 mg/ml) were adsorbed on carbon formvar-coated copper grids and were negatively stained with 0.5% uranyl acetate aqueous solution. The grids were examined using a JEM-1230 electron microscope (JEOL, Tokyo, Japan) at an accelerating voltage of 100 kV.

### Mass Spectrometry

Wild type CCMV_TT_-VLPs, Round or Rod-shaped CCMV_TT_-VLPs (1 mg/ml in buffer A) were diluted with a 2,5-Dihydroxyacetophenone (2,5-DHAPI) matrix solution and were spotted onto an MTP AnchorChip 400/384TF. Matrix assisted laser desorption/ionization (MALDI)-TOF MS analysis was carried out on an Autoflex MS (Bruker Scientific, Billerica, Massachusetts). The protein molecular mass (MM) calibration standard II (22.3–66.5 kDa; Bruker, Billerica, Massachusetts) was used for mass determination.

### SDS-Page and gel electrophoresis

SDS-Page: 6µg of Round or Rod-shaped CCMV_TT_-VLPs were mixed with 2x mercaptoethanol and heated at 95°C for 3 minutes and then loaded into Any kD Mini-PROTEAN TGX precast protein gels (BIO-RAD). Gel was run for 35min at 180V. As reference Page Ruler™ Prestained Protein Ladder was used (Thermo Fisher Scientific, Waltham, Massachusetts). Gel electrophoresis: 10µg of Round- or Rod-shaped CCMV_TT_-VLPs were loaded on a 1% agarose gel. Nucleic acids were visualized using Cybr Safe DNA Gel Stain (Life Technologies, Carlsbad, California). 5µl Quick-Load Purple 1 kb DNA ladder (New England Biolabs, Ipswich, Massachusetts) was used as reference. Gel was run for 30min at 100V.

### Mice

Wild type C57BL/6 mice were purchased from Harlan. All *in vivo* experiments were performed using 8-12 weeks old female mice. All animal procedures were conducted in accordance with the Swiss Animal Act (455.109.1 – September 2008, 5^th^).

### Vaccination regimen

Wild type C57BL/6 mice (8-12 weeks, Harlan) were vaccinated subcutaneously (SC) with 15μg Round or Rod-shaped CCMV_TT_-VLPs in 100µl PBS on day 0. Mice were boosted with a identical dose on day 14. Serum samples were collected on days 0, 7, 14, 21, 28 and 35.

### The enzyme-linked immunosorbent assay (ELISA)

For determination of total IgG antibody titers against Round and Rod-shaped CCMV_TT_-VLPs in sera of immunized mice, ELISA plates (Nunc Immuno MaxiSorp, Rochester, NY) were coated over night with Round or Rod-shaped CCMV_TT_-VLPs, respectively. Plates were washed with PBS-0.01%Tween and blocked using 100µl PBS-Casein 0.15% for 2h. Sera from immunized mice were diluted 1/20 initially and a 1/3 dilution chain was performed. Plates were incubated for 1h at 37°C. After washing with PBS-0.01%Tween, goat anti-mouse IgG conjugated to Horseradish Peroxidase (HRP) (Jackson ImmunoResearch, West Grove, Pennsylvania) was added 1/1000 and incubated for 1h at 37°C. Plates were developed and OD 450 reading was performed.

IgG subclasses were measured from day 21 sera using the same ELISA protocol with the following secondary Abs: goat anti-mouse IgG1-HRP (BD Biosciences, San Jose, California), goat anti-mouse IgG2b-HRP (1:1000) (Thermo Fischer Scientific, Waltham, Massachusetts), goat anti-mouse IgG2c-HRP (Southern Biotech, Birmingham, Alabama) 1:4000, rat anti-mouse IgG3-biotin (Becton, Dickinson, Franklin Lakes, New Jersey) 1:2000 followed by streptavidin-HRP (Dako, Glostrup, Denmark) 1:1000 incubated at 37°C for 1h.

IgA was measured using day 35 sera (immunization at day 0, boost at day 14). IgG was depleted using Dynabeads™ Protein G (Thermo Fischer Scientific, Waltham, Massachusetts). Serum was diluted 1/20 (total volume of 75µl) in PBS-Casein 0.15%. 25µl beads were used per sample. Manufacturer’s protocol was followed until step 3. of “Bind Antibody”. Supernatant was added to ELISA plates and a 1/3 serial dilution was performed. For IgA detection, goat anti mouse IgA conjugated to HRP was used (ThermoFisher Scientific, Waltham, Massachusetts) 1/4000.

For OD50 calculations, if a sample did not reach the threshold a value of 1 was appointed.

### Measuring IFN-γ in mouse serum

Wild type C57BL/6 mice (8-12 weeks, Harlan) were vaccinated with 15µg Round or Rod-shaped CCMV_TT_-VLPs on day 0 and boosted on day 14. Serum from vaccinated mice was collected for measuring IFN-γ. ELISA MAX™ Deluxe Set Mouse IFN-γ (Biolegend, San Diego, California) was performed according to manufacturer’s instructions. Serum was used undiluted and concentration was interpolated to a standard curve of the sets standard sample.

### Trafficking of Round and Rod-shaped CCMV_TT_-VLPs to draining lymph nodes

Round or Rod-shaped CCMV_TT_-VLPs were labelled with AF488 as per manufacturer’s instructions (Thermo Fischer Scientific, Waltham, Massachusetts) and stored at -20°C. Wild type C57BL/6 mice (8-12 weeks, Harlan) were injected with 10µg of the VLPs in the footpad under isoflurane anesthesia. Popliteal LNs were collected 3h and 24h following footpad injection. Lymph nodes were treated with collagenase D (Roche, Basel, Switzerland) and DNase I (Boehringer, Ingelheim am Rheih, Germany) in DMEM medium containing 5% FBS and 1% Strep/Penicillin for 30 min at 37°C. Lymph nodes were smashed using 70µm cell strainers, RBC were lysed with ACK buffer. Cells were stained with Fc blocker and then with anti-CD11b, CD11c, CD45R/220, CD8 and F4/80 (all from Biolegend, San Diego, California).

### Immunofluorescence

Round or Rod-shaped CCMV_TT_-VLPs were labelled with AF488 as per manufacturer’s instructions (Thermo Fischer Scientific, Waltham, Massachusetts) and stored at -20°C. Wild type C57BL/6 mice (8-12 weeks, Harlan) were injected with 10µg of the VLPs in the footpad under isoflurane anesthesia. Popliteal LNs were collected 3h and 24h following footpad injection and embedded in Tissue-Tek optimum cutting temperature compound (Sakura) without prior fixation. Cryostat sections (7μm in thickness) on Superforst/Plus glass slides (Thermo Fischer Scientific, Waltham, Massachusetts) were air-dried overnight and then fixed for 10 min in ice-cold 100% acetone (PanReac). After rehydration (5 min in 1x PBS), sections were blocked with 1% (w/v) BSA (Sigma Aldrich, St. Louis, Missouri) and 1% (v/v) normal mouse serum. Immunofluorescence labeling was done with Abs diluted in PBS containing 0.1% (w/v) BSA and 1% (v/v) normal mouse serum. Sections were washed 3 times for 5 min in 1x PBS after every labeling step. LN staining: macrophages were detected using a primary antibody against CD11b (1/1000, rat anti mouse CD11b conjugated with PE; BD Biosciences, San Jose, California), B-cell follicles were identified using rat anti mouse B220 Alexa F647 (1/1000; BD Biosciences, San Jose, California). Images were acquired on an Axioplan microscope using an AxioCam MRn (Zeiss).

### Histology of lymph node

Round or Rod-shaped CCMV_TT_-VLPs were labelled with AF488 as per manufacturer’s instructions (Thermo Fischer Scientific, Waltham, Massachusetts) and stored at -20. Wild type C57BL/6 mice (8-12 weeks, Harlan) were injected with 10µg of the VLPs in the footpad under isoflurane anesthesia. Popliteal LNs were collected 3h and 24h following footpad injection and fixed with 4% paraformaldehyde solution (Sigma Aldrich, St. Louis, Missouri). Of each group, two to four murine popliteal LNs were histologically examined by a board-certified veterinary pathologist (SdB). Of each LN, a full cross section, stained with Hematoxylin and Eosin (HE), was assessed for any histopathological changes.

### Statistical analysis

Data were analyzed and presented as mean ± SEM using GraphPad PRISM 8. Comparison between the groups was performed using Student’s *t*-test. *P*-values ^****^*P* < 0.0001; ^***^*P* < 0.001; ^**^*P* < 0.01; ^*^*P* < 0.05.

## Results

### Directional insertion of tetanus toxoid (TT) epitope in the N or C-terminal results in Round or Rod-shaped CCMV_TT_-VLPs

In a first step, we engineered CCMV-VLPs derived from a non-human pathogenic plant virus by incorporating a powerful T-cell stimulatory epitope derived from tetanus toxoid (TT) (Tetanus toxin 830-843) at the N or C-terminus of CCMV-VLPs. The TT epitope was genetically fused to the capsid protein (CP) of CCMV-VLPs as has been previously described for our newly developed platform derived from cucumber-mosaic virus-like particles (CuMV_TT_-VLPs) (31). The introduced TT epitope is considered a universal epitope in humans as it is recognized by essentially all individuals. Since all individuals have been immunized against TT, memory Th cells may be able to help B cells to generate protective IgG even under more challenging conditions such as aged populations (31). CCMV-VLPs with insertion of TT epitope at the N-terminus retain their self-assembly as an icosahedron similar to the native virus. In contrast, insertion of TT epitope at the C-terminal end led to formation of Rod-shaped particles. Both Round and Rod-shaped CCMV-VLPs forms carry a lysine to arginine mutation at residue 42 (32). This mutation renders the VLPs less sensitive to pH and salt concentration, which is an advantage in *in vivo* environments (A. Zeltins, manuscript in preparation). Therefore, the engineered VLPs in this study are salt-stable (SS). The shape and integrity of the cloned VLPs were confirmed via electron microscopy which shows a size of ∼30nm in diameter for CCMV-Ntt-SS (Fig. 1a). The size of the CCMV_TT_-Ctt-SS greatly varies in length but can reach up to more than 1µm, with a width of about 30nm. Figure 1b shows a magnified image of CCMV_TT_-Ctt-SS VLPs for easy comparison of their width with the icosahedral CCMV_TT_-Ntt-SS VLPs in figure 1a. To reach a rather homogenous population, we performed sucrose gradient separation to focus on the long rods (Fig. 1c). For simplification we refer to the two forms of engineered CCMV_TT_ in this paper as Round-shaped CCMV_TT_-VLPs (CCMV-Ntt-SS) and Rod-shaped CCMV_TT_-VLPs (CCMV-Ctt-SS).

**Figure 1:**
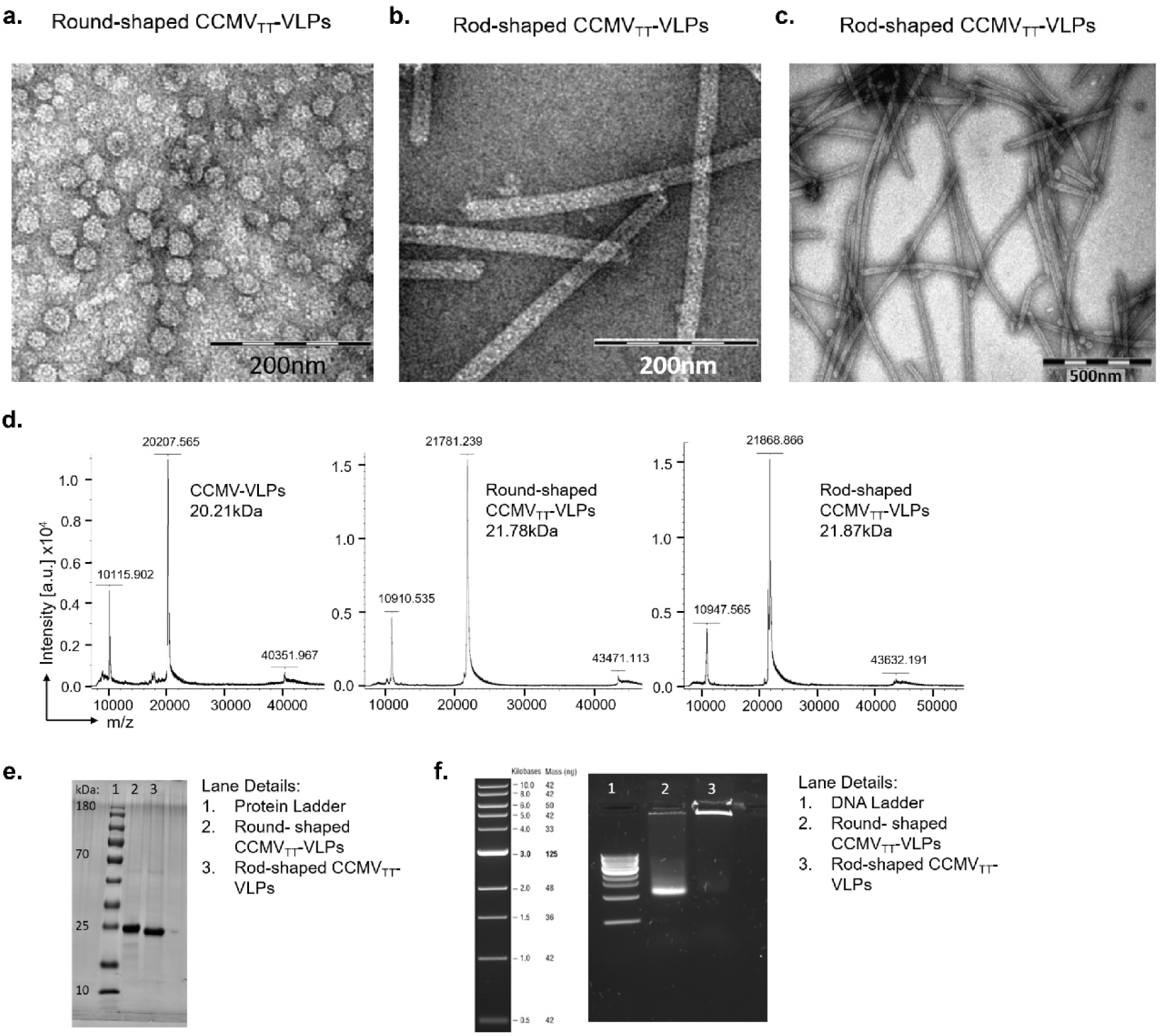
Directional insertion of tetanus toxoid (TT) epitope in the N or C-terminal results in Round or Rod-shaped CCMV_TT_-VLPs. ***a***, EM of Round-shaped and ***b***.***/c.*** Rod-shaped CCMV_TT_-VLPs, adsorbed on carbon grids and negatively stained with uranyl acetate solution, scale bars 200nm (Round) and 200nm/500nm (Rod). Round-shaped CCMV_TT_-VLPs are ∼30nm in diameter, Rod-shaped CCMV_TT_-VLPs are ∼1µm in length and ∼30nm in width. ***d***, Mass spectrometry data, from left to right: wild type CCMV_TT_-VLPs, Round-shaped CCMV_TT_-VLPs and Rod-shaped CCMV_TT_-VLPs. ***e***, Reducing SDS-Page stained with Coomassie-blue stain, lane 1: protein marker, lane 2: Round-shaped CCMV_TT_, lane 3: Rod-shaped CCMV_TT_. ***f***, Agarose gel stained with SYBR safe, lane 1: DNA ladder, lane 2: Round-shaped CCMV_TT_, lane 3: Rod-shaped CCMV_TT_.

To further characterize the two forms of CCMV_TT_, we performed mass spectrometry (MS). MS data revealed a molecular weight for the CP monomers of Round- and Rod-shaped CCMV_TT_-VLPs of roughly 21.8 and 21.9kDa, respectively (Fig. 1d). The original CCMV-SS (salt-stable CCMV without TT insertion) is formed by CPs of roughly 20.2kDa. Considering the weight of the TT 830-843 epitope of 1.611kDa the obtained data are consistent. Reducing SDS-page experiments confirmed these findings and showed bands for Round- and Rod-shaped CCMV_TT_-VLPs of equal height at the appropriate position (Fig. 1e).

Both engineered CCMV_TT_-VLPs were produced in an *E*. coli system. The VLPs packaged ssRNA derived from *E*. coli spontaneously, which serves as a potent TLR7/8 ligand. The concentration of RNA in both Round- and Rod-shaped CCMV_TT_-VLPs was similar when measured at 260nm. The RNA content of the Round-shaped CCMV_TT_-VLPs could also be visualized by agarose gel electrophoresis, however this was less efficient for the Rod-shaped CCMV_TT_-VLPs because most of the RNA staying in the slot based on the Rod-shaped CCMV_TT_-VLPs being unable to move through the gel (Fig. 1f).

### Round-shaped CCMV_TT_-VLPs exhibit faster and more efficient draining kinetics than Rod-shaped CCMV_TT_-VLPs in vivo

To test the role of size and shape in lymphatic trafficking of the engineered VLPs, we assessed and visualized the accumulation of the AF488 labelled Round or Rod-shaped CCMV_TT_-VLPs in murine popliteal lymph nodes (LNs) 3h and 24h after injection in the footpads. Flow cytometry data confirms that Round-shaped CCMV_TT_-VLPs were more efficient in reaching the popliteal LNs than the Rod-shaped ones. Such finding is not surprising as the round VLPs have a size of ∼30nm in diameter allowing them to drain freely through the 200nm pores of the lymphatic vessels. Nevertheless, the Rod-shaped CCMV_TT_-VLPs were also capable of draining through the lymphatic vessels within 3 hours, despite their micro-size, this may be due to the fact that the width of the rods is ∼30nm, similar to the diameter of the round-shaped CCMV_TT_-VLPs.

Next, we studied which type of APCs are involved in interacting with the Round and Rod-shaped CCMV_TT_-VLPs. Lymph node conventional dendritic cells (cDCs); CD8^+^CD11c^+^ (Fig. 2a), CD8^-^CD11c^+^ (Fig. 2b), macrophages CD11b^+^F4/80^+^ (Fig. 2c), different myeloid cells populations CD45^+^CD11b^+^ (Fig. 2d and h), neutrophils CD45^+^ Ly6G^+^ (Fig. 2e) and B cells CD45R/B220^+^ (Fig. 2f and i) were more efficient in taking up Round-shaped than Rod-shaped CCMV_TT_-VLPs 3h after injection. No significant difference in the uptake between Round and Rod-shaped CCMV_TT_-VLPs has been seen at 24h except for B cells and the macrophage population characterized by CD11b^+^F4/80^+^ (Fig. 2c). Interestingly, uptake by those macrophages was more prominent for Rod-shaped than Round-shaped CCMV_TT_-VLPs 24h post injection. This might result from delayed drainage of Rod-shaped CCMV_TT_-VLPs. B cells characterized by CD45R/B220^+^ showed a very high frequency in taking up Round-shaped CCMV_TT_-VLPs in the popliteal LNs 3h and 24h after injection in the mouse footpad. B cells taking up CCMV_TT_-VLPs also showed expression of MHC-II (Fig. 2f and g). We next followed the arrival of the labelled CCMV_TT_-VLPs to the draining LNs by fluorescence microscopy of cryosections obtained from excised popliteal LNS at 3h and 24h post injection in the mouse footpad. The sections were co-stained with macrophage and B-cell markers, CD11b^+^ and CD45^+^/B220^+^, respectively. The results demonstrate that AF488 Round-shaped CCMV_TT_-VLPs accumulated in large numbers in the subcapsular sinus area (SCS), the cortex and the medullary sinus (MS) of the popliteal LNs 3h post injection (Fig. 2j). Macrophages characterized with CD11b^+^ were prominent 3h post injection with Round-shaped CCMV_TT_-VLPs at the SCS and MS. Rod-shaped CCMV_TT_-VLPs were less visible 3h at the popliteal LN post injection in the footpad and their presence was confined to the SCS with scarce VLPs in the cortex. 24h post injection, Round-shaped CCMV_TT_-VLPs were found deeper in the popliteal LN. Whereas, this observation was less obvious for the Rod-shaped CCMV_TT_-VLPs. B-cells were detectable 3h and 24h post injection of the VLPs in the mouse footpads. As the accumulation of Round-shaped CCMV_TT_-VLPs was prominent 3h post injection in the footpad, the B-cell signal was less noticeable. 24h later the accumulation of B-cells was more pronounced at the SCS and the cortical area of the LN upon injection of Round-shaped CCMV_TT_-VLPs than after injection with Rod-shaped CCMV_TT_-VLPs.

**Figure 2:**
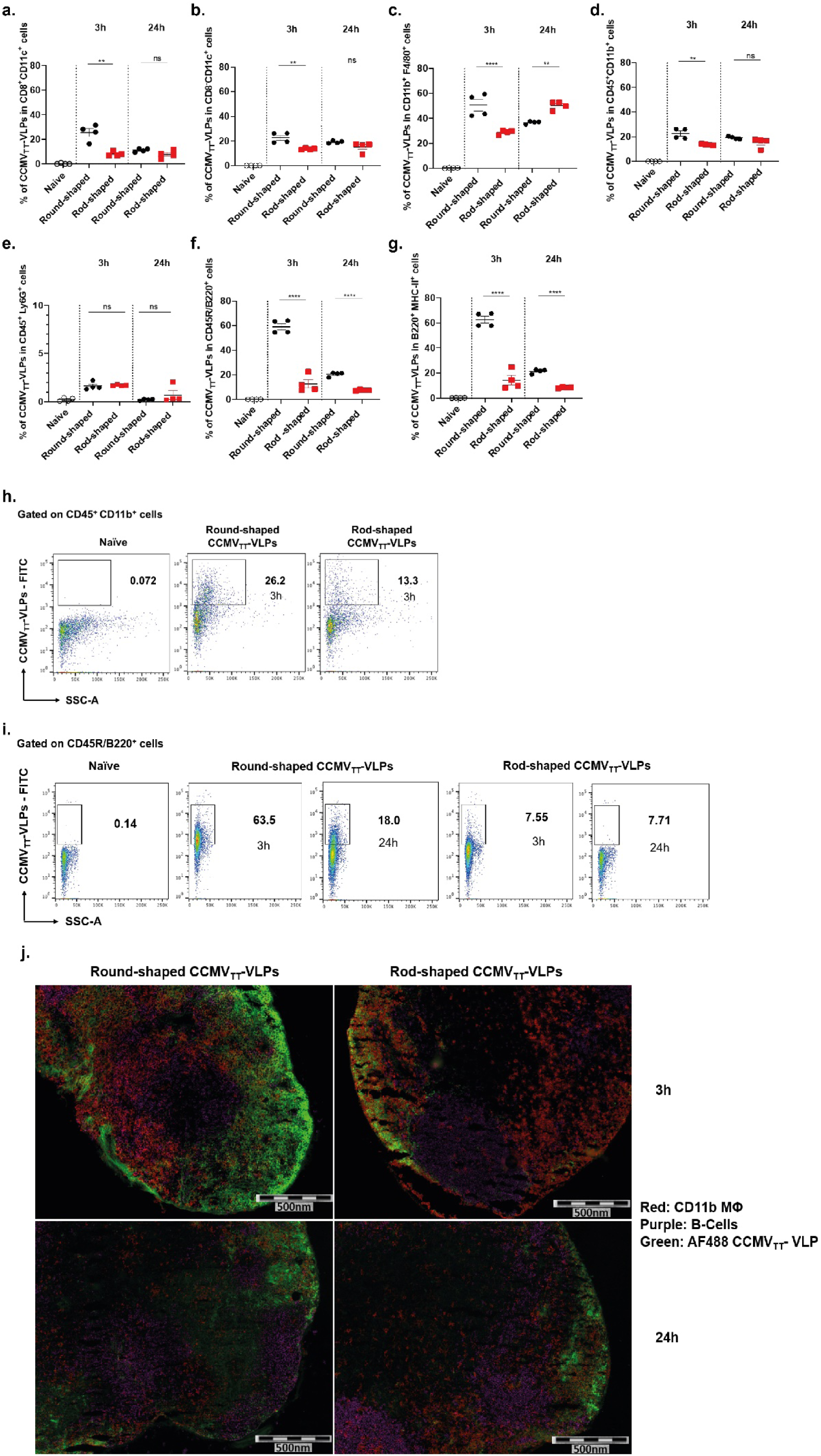
Round-shaped CCMV_TT_-VLPs exhibit faster and more efficient draining kinetics than Rod-shaped CCMV_TT_-VLPs in vivo. Percentage of different cell populations positive for AF488 labeled Round and Rod-shaped CCMV_TT_-VLPs in murine popliteal LNs 3h and 24h post injection in footpad: ***a***, CD8^+^CD11c^+^ cells; ***b***, CD8^-^CD11c^+^ cells; ***c***, CD11b^+^F4/80^+^ cells; ***d***, CD45^+^CD11b^+^ cells. ***e***, CD45^+^Ly6G^+^ cells; ***f***, CD45R/B220^+^ cells and ***g***, CD45R/B220^+^ MHC^+^ cells. ***h***, Representative FACS plots showing the percentage of CCMV_TT_-VLPs labelled with AF488 in CD45^+^CD11b^+^ cells in popliteal LNs 3h post injection in mouse footpad. ***i***, Representative FACS plot showing the percentage of CCMV_TT_-VLPs labelled with AF488 in CD45R/B220^+^ cells in popliteal LNs 3h and 24h post injection in mouse footpad. Naïve LNs were used as a negative control. ***j***, Immunofluorescence of popliteal LNs 3h and 24h post vaccination with Round or Rod-shaped CCMV_TT_-VLPs labeled with AF488, cyrosections were treated with Abs detecting CD11b^+^ cells (red colour) and CD45/B220^+^ cells (purple colour). Mean ± SEM, 4 mice per group, one representative of 2 similar experiments is shown.

### Round-shaped CCMV_TT_-VLPs are more potent in inducing IgG antibodies than Rod-shaped CCMV_TT_-VLPs

In a next step, we have assessed the humoral immune response induced by both engineered Round and Rod-shaped CCMV_TT_-VLPs. It is important to keep in mind that it is usually difficult to compare different sized particles for their immunogenicity as for spheres, the surface is proportional to radius (r)^2^, while the weight will be proportional to (r)^3^. Hence the weight of the injected particles grows much more rapidly than the surface, rendering a comparison difficult. In contrast, for rods, both the surface and weight are proportional to the length of the rod, rendering this comparison more appropriate.

Hence, C57BL/6 mice were vaccinated subcutaneously (SC) with 15µg of Round or Rod-shaped CCMV_TT_-VLPs on day 0 and boosted once on day 14 as illustrated in Figure 3a. Serum was collected on day 0 before vaccination and subsequently on day 7, 14, 21, 28 and 35. Total specific IgG response against Round or Rod-shaped CCMV_TT_-VLPs was assessed by ELISA. As shown in Fig. 3b and c, the Round-shaped CCMV_TT_-VLPs were very potent at inducing specific IgG response 7 days following the administration of the first dose. This response was enhanced dramatically on days 14, 21, 28 and 35. On the contrary, vaccination with Rod-shaped CCMV_TT_-VLPs was low after the first dose and has been increasing significantly only following boost on day 14.

**Figure 3:**
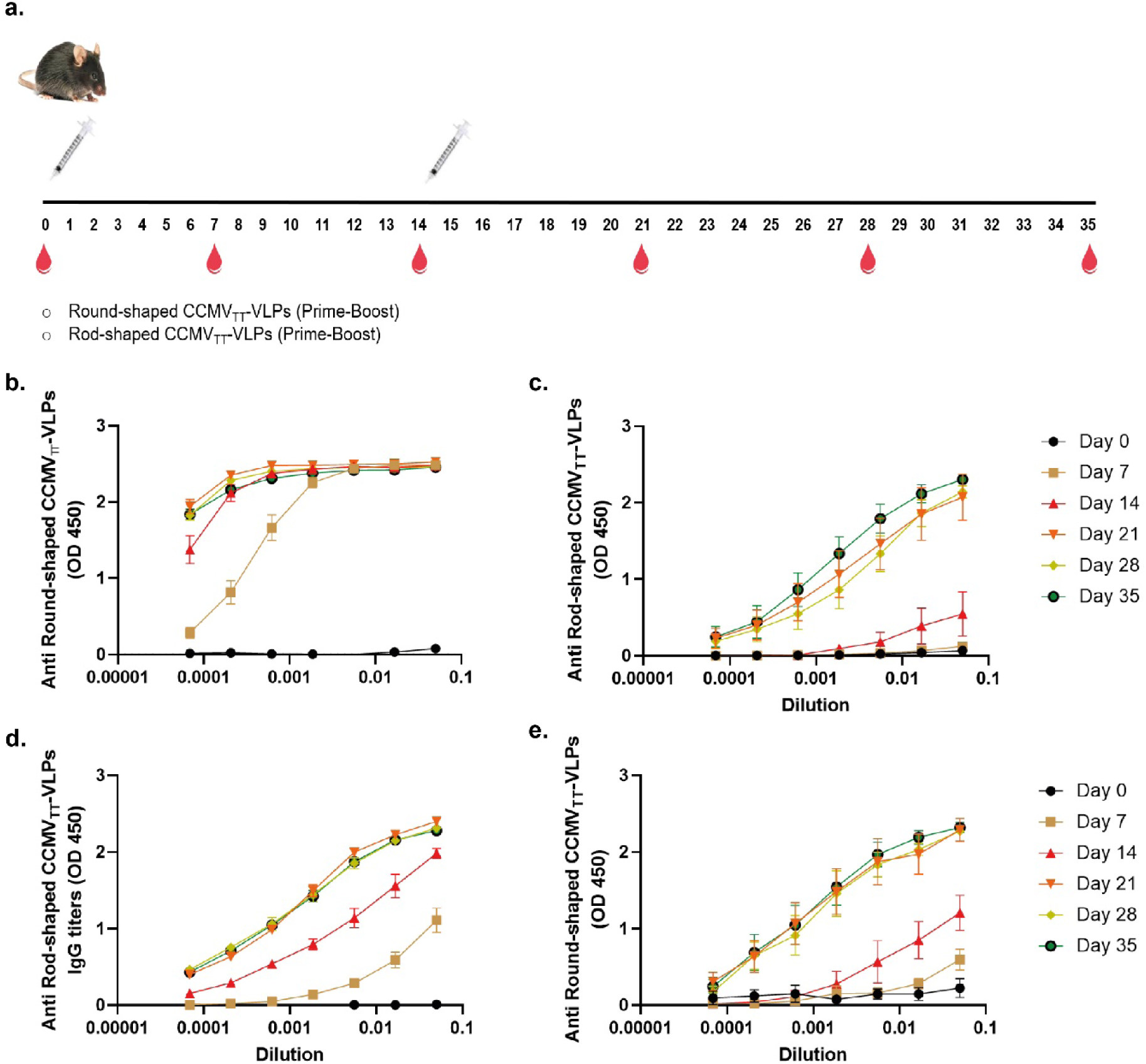
Round-shaped CCMV_TT_-VLPs are more potent in inducing IgG antibodies than Rod-shaped CCMV_TT_-VLPs. ***a***, Vaccination regimen and bleeding time-points. ***b***, Sera from Round-shaped CCMV_TT_-VLP vaccinated mice, ELISA plates coated with Round-shaped CCMV_TT_-VLPs. ***c***, Sera from Rod-shaped CCMV_TT_-VLP vaccinated mice, ELISA plates coated with Rod-shaped CCMV_TT_-VLPs. ***d***, Sera from Round-shaped CCMV_TT_-VLP vaccinated mice, ELISA plates coated with Rod-shaped CCMV_TT_-VLPs. ***e***, Sera from Rod-shaped CCMV_TT_-VLP vaccinated mice, ELISA plates coated with Round-shaped CCMV_TT_-VLPs. Mean ± SEM, 6 mice per group, one representative of 2 similar experiments is shown.

Even though subunits of both VLPs were almost identical, in order to test the cross-reactivity of both Round and Rod-shaped CCMV_TT_-VLPs, we have tested the collected sera in ELISA coated with the opposite VLP shape. Specifically, sera from mice vaccinated with Round-shaped CCMV_TT_-VLPs were tested against ELISA coated with Rod-shaped CCMV_TT_-VLPs and vice versa. Our results showed that sera from mice vaccinated with Round-shaped CCMV_TT_-VLPs are capable of recognizing the Rod-shaped CCMV_TT_-VLPs after a single dose on day 7. The response was enhanced on day 14, 21, 28 and 35. However, sera from mice vaccinated with Rod-shaped CCMV_TT_-VLPs could only significantly recognize the Round-shaped CCMV_TT_-VLPs after boosting (Fig. 3d and e). In a next step, we have produced Rod-shaped CCMV_TT_-VLPs exhibiting variation in lengths that includes smaller fragmented pieces of less than ∼1µm in length (By ways of using polyethylene glycol precipitation instead of sucrose gradient centrifugation for purification, as described for Round-shaped CCMV_TT_-VLPs in material & methods; Supplementary Fig. 1). Specific anti-IgG response against the Rod-shaped

CCMV_TT_-VLPs was enhanced substantially, indicating that indeed the rod-length limited the size of the immune response (Supplementary Fig. 2). In all other experiments presented in this paper the more homogeneous long Rod-shaped CCMV_TT_-VLPs were used.

### Rod-shaped CCMV_TT_-VLPs fail to induce isotype switching in comparison to the Round-shaped CCMV_TT_-VLPs

The C57BL/6 murine IgG family consists of four major subclasses IgG1, IgG2b, IgG2c and IgG3. Each unique subclass is implicated in distinct effector functions during humoral immune responses. VLPs and other nanoparticles induce a humoral response dominated by IgG1 in the absence of packaged RNA or DNA in mice (26, 33, 34). Isotype switching to the protective IgG2 subtype is strictly TLR dependent. Thus, VLPs packaged with prokaryotic ssRNA during *E*. coli production induce a humoral immune response dominated with IgG2 subclasses (35, 36). Based on these grounds, we were interested in characterizing the different IgG subclasses in mice sera vaccinated with Round vs Rod-shaped CCMV_TT_-VLPs. As explained earlier the engineered Round and Rod-shaped CCMV_TT_-VLPs package the same quantity of ssRNA. Therefore, TLR7/8 ligand effect can be eliminated as a confounding variable and the difference in the induced IgG subclasses can be correlated to the size of the VLPs. Our analysis revealed that Round-shaped CCMV_TT_-VLPs significantly enhanced all IgG subclasses compared to the Rod-shaped ones. The difference, however, was most striking for TLR7/8 related subclasses IgG2b/c and IgG3 (Fig.4a). By performing OD50 analysis we compared the titers (given as reciprocal dilution values) in both groups as depicted in Figure 4b. Titers are significantly higher (*p*. <0.0001 for IgG1, IgG2b and IgG2c) (p. <0.05 for IgG3) post immunization with Round-shaped CCMV_TT_-VLPs. The ratio between TH_1_ and TH_2_ associated IgG subclasses was calculated next and it became evident that TH_1_ contribution is more pronounced with Round-shaped CCMV_TT_-VLPs (Fig. 4c).

**Figure 4:**
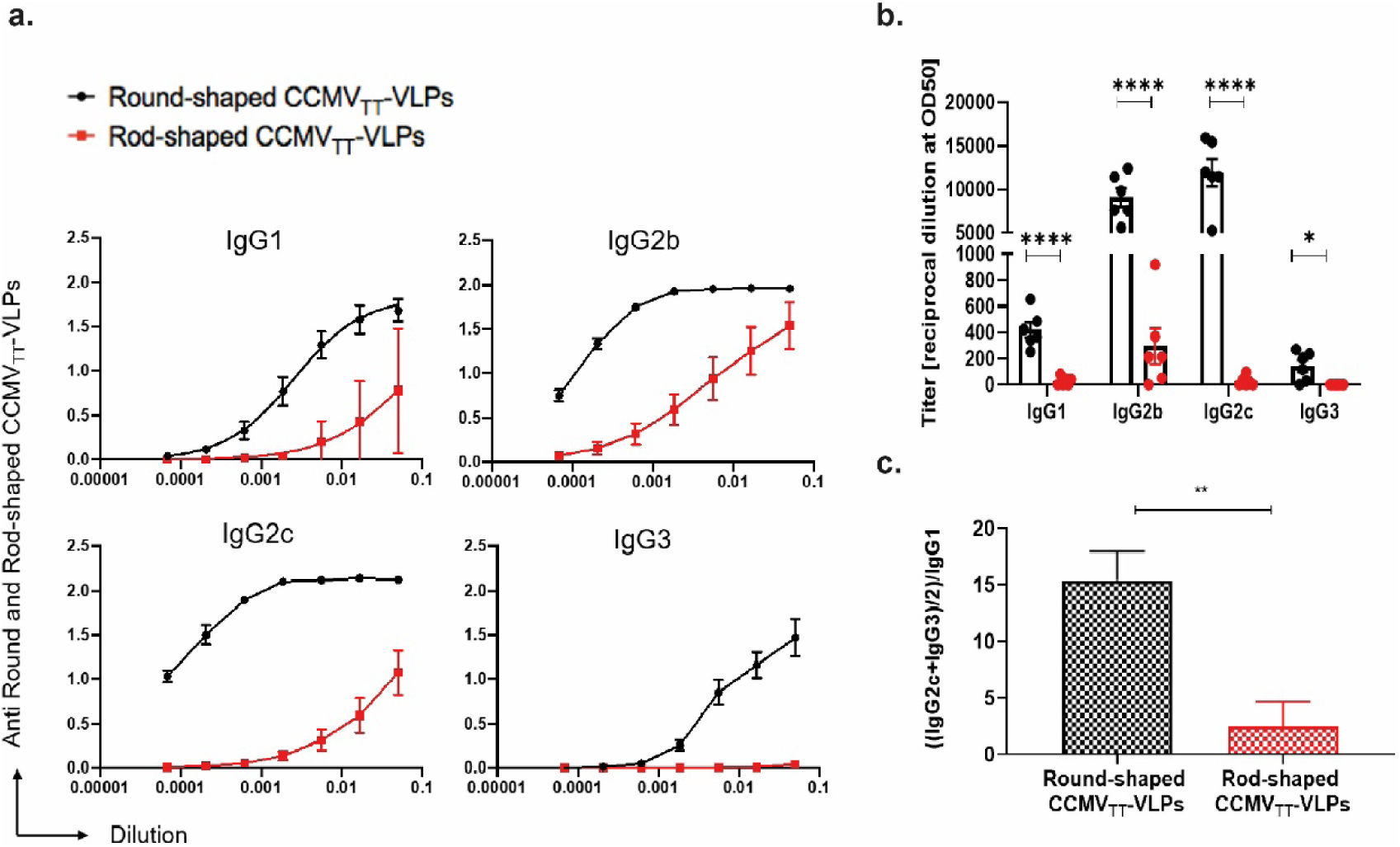
Rod-shaped CCMV_TT_-VLPs fail to induce isotype switching in comparison to the Round-shaped CCMV_TT_-VLPs. ***a***, Anti-Round and Rod-shaped CCMV_TT_-VLP specific IgG1, IgG2b, IgG2c and IgG3 titers measured in day 21 mice sera using OD450nm. ELISA plates were coated with Round-shaped CCMV_TT_-VLPs for detecting IgG subclasses in mice vaccinated with Round-shaped CCMV_TT_-VLPs. ELISA plates were coated with Rod-shaped CCMV_TT_-VLPs for detecting IgG subclasses in mice vaccinated with Rod-shaped CCMV_TT_-VLPs. ***b***, Anti-Round and Rod-shaped CCMV_TT_-VLP specific IgG1, IgG2b, IgG2c and IgG3 titers measured in day 21 mice sera using OD50 calculation of data depicted in a. ***c***, TH_1_/TH_2_ ratio in Round and Rod-shaped CCMV_TT_-VLP immunized mice depicted as ((IgG2c+IgG3)/2)/IgG1 of OD50 values shown in b. Mean ± SEM, 6 mice per group, one representative of 2 similar experiments is shown.

### Systemic IgA response depends on size of VLPs

Previous studies have shown that SC injection of VLPs packaging RNA leads to a strong IgA response despite the fact that IgA antibodies are TH-cell independent (35). Besides, our previous findings revealed that systemic IgA response is heavily dependent on TLR7/8 in B-cells (37). The role of the size of VLPs packaging similar contents of RNA has not been investigated before, therefore we carried out an experiment to investigate this matter. Our findings indicate that Round-shaped CCMV_TT_-VLPs could induce significantly higher (*p*. <0.0001) isotype switching to IgA when compared to the Rod-shaped ones (Fig. 5a and b). To rule out the role of TH-cells, we have also measured IFN-γ in the serum of vaccinated mice on day 14, the results showed no significant difference (*p*. 0.8211) between both groups (Fig. 5c).

**Figure 5.**
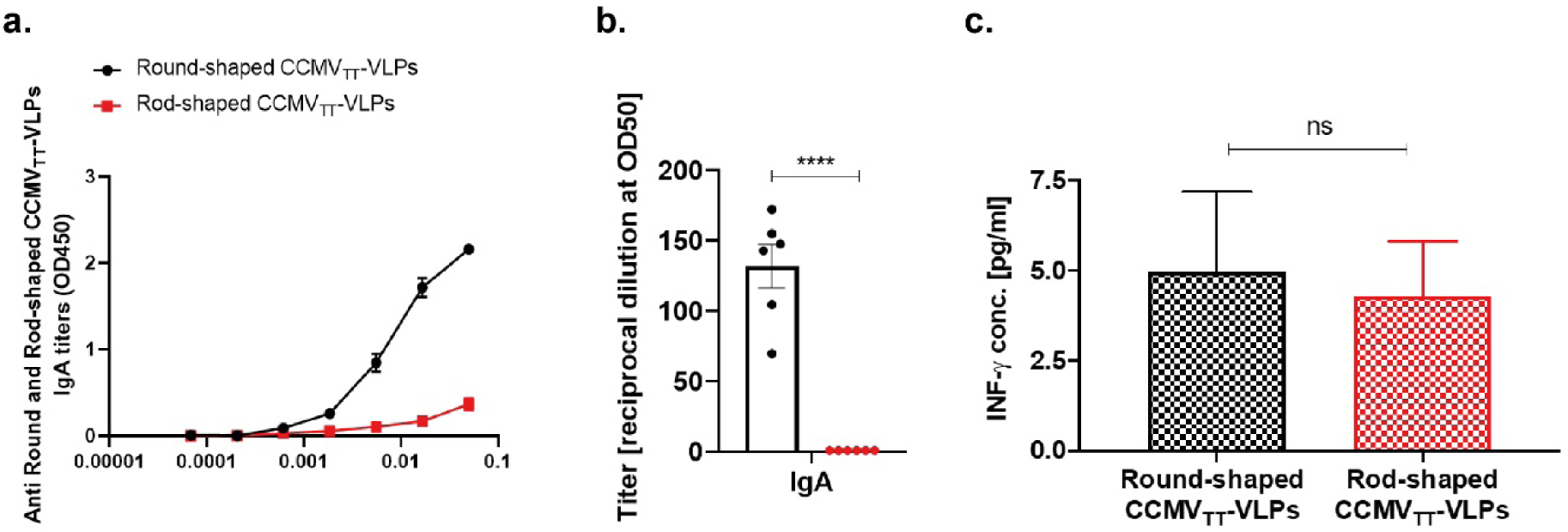
Systemic IgA response depends on size of VLPs. ***a***, Anti-Round and Rod-shaped CCMV_TT_-VLP specific IgA titers measured in day 21 serum from mice vaccinated with Round and Rod-shaped CCMV_TT_-VLPs using OD450nm. ***b***, Anti-Round and Rod-shaped CCMV_TT_-VLP specific IgA titers measured using OD50 calculation of data depicted in a. ***c***, Concentration of IFN-γ measured in day 14 serum from mice vaccinated with Round and Rod-shaped CCMV_TT_-VLPs. Mean ± SEM, 6 mice per group, one representative of 2 similar experiments is shown.

### Germinal center formation is prominent following a single dose of Round-shaped CCMV_TT_-VLPs

Antigen reservoir persisting on follicular dendritic cells (fDCs) is essential for germinal centers (GC) to keep B-cells stimulated and to generate a strong and long-lived antibody response. We therefore studied the formation of GCs in the spleens of mice vaccinated with Round and Rod-shaped CCMV_TT_-VLPs 12 days following a single SC dose of the engineered VLPs. Results showed that the formation of GCs in mice vaccinated with Round-shaped CCMV_TT_-VLPs was significantly higher than in Rod-shaped CCMV_TT_-VLP vaccinated mice when measuring the total number of GCs (*p*. 0.0042) or the number of GCs/mm^2^ (*p*. 0.0033) (Fig. 6a and b). Histological examination of Hematoxylin and Eosin (HE) stained splenic tissue indicated the presence of multiple GCs within the lymphoid follicles upon vaccination with Round-shaped CCMV_TT_-VLPs. On the other hand, mice vaccinated with Rod-shaped CCMV_TT_-VLPs revealed only rare GC formation. Spleens from naïve C57BL/6 mice were used as a control (Fig. 6c and d). The examined spleens in mice vaccinated with Round or Rod-shaped CCMV_TT_-VLPs were free of any relevant degenerative or necrotic histopathological changes.

**Figure 6:**
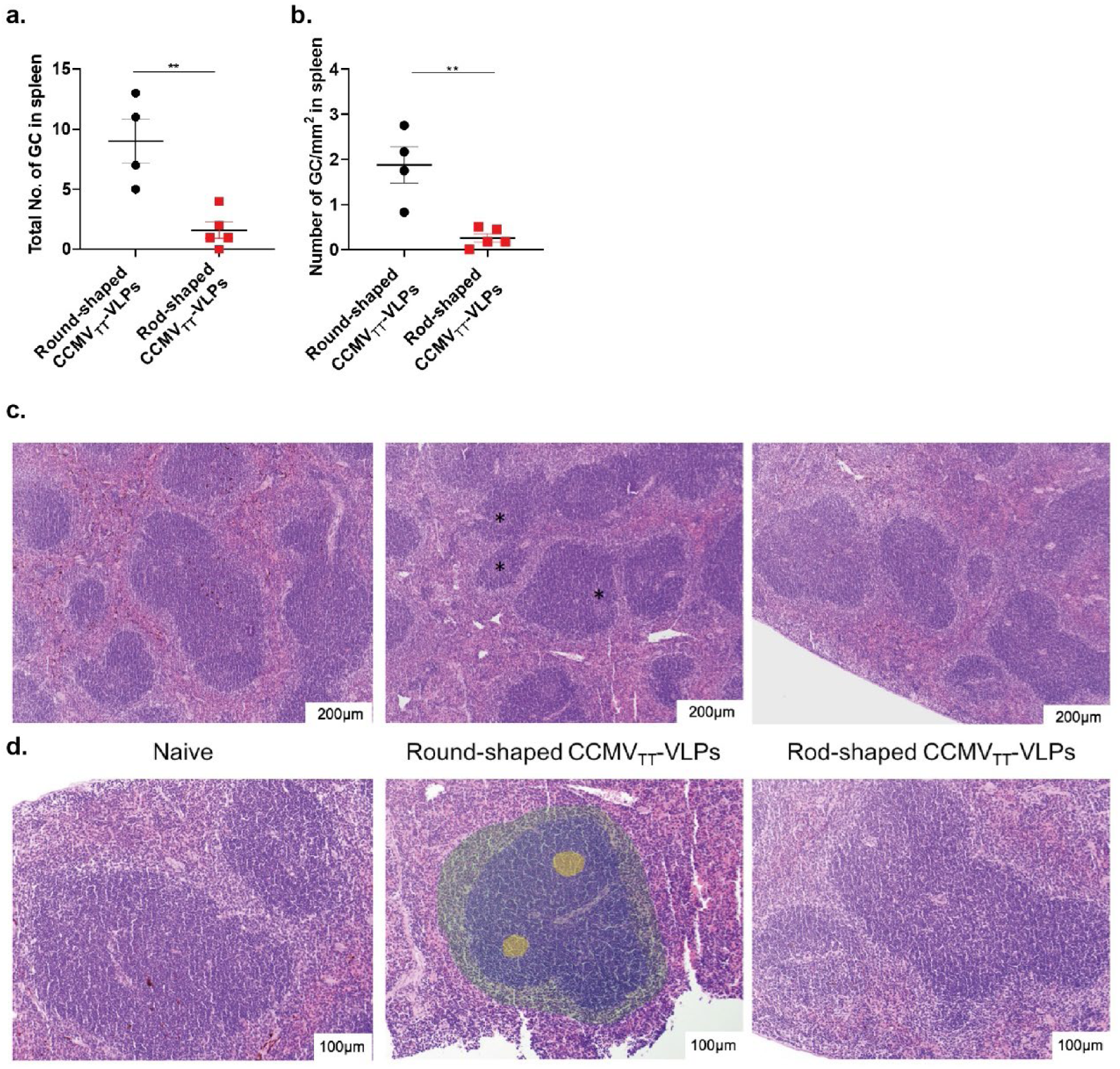
Germinal center formation is prominent following a single dose of Round-shaped CCMV_TT_-VLPs. ***a***, Total number of germinal centers (GCs) in the examined splenic tissue fragments of mice vaccinated with a single dose of 15μg of Round or Rod-shaped CCMV_TT_-VLPs. ***b***, Number of GCs/mm^2^ in spleen of mice vaccinated with a single dose of 15μg of Round-shaped CCMV_TT_-VLPs. ***c***, Histology of HE stained murine spleens, 10x objective. From the left: naïve spleen, lymph follicles lack evident GC formation; spleen of mice vaccinated with Round-shaped CCMV_TT_-VLPs, lymph follicle present multifocal GC formation (*); last spleen of mice vaccinated with Rod-shaped CCMV_TT_-VLPs, lymph follicle lack visible GCs. ***d***, Higher magnification of c., 20x objective. From the left: naïve spleen, lymph follicles present with densely arranged lymphocytes without evident GC formation; spleen of mice vaccinated with Round-shaped CCMV_TT_-VLPs, GCs (*) are visible within the lymph follicles, GCs and surrounding follicular and perifollicular zones are highlighted with colours (GCs (yellow), mantle zone (blue) and marginal zone (green)); last spleen of mice vaccinated with Rod-shaped CCMV_TT_-VLPs, lymph follicles present with densely arranged lymphocytes without evident GC formation. Mean ± SEM, 2 splenic tissue fragments were analyzed from naïve group, 4 from Round-shaped CCMV_TT_-VLPs group and 5 from Rod-shaped CCMV_TT_-VLPs group.

### Round and Rod-shaped CCMV_TT_-VLPs enhance neutrophil infiltration in lymph node sinuses without causing any degenerative or vascular changes

To study the histopathological effect of Round and Rod-shaped CCMV_TT_-VLPs on LNs, we have performed histology of HE stained popliteal LNs collected 3h and 24h following footpad injections. The main histopathological change of the examined LNs was restricted to a mild to moderate increase of neutrophils within the sinuses, which was present in both groups and absent or neglectable in control tissue (Table 1 and Fig. 7a-c). This neutrophil infiltrate was more evident 3h than 24h after injection. No convincing histological difference was observed between the two groups vaccinated with Round or Rod-shaped CCMV_TT_-VLPs. Evidence of tissue damage, i.e. degeneration or necrosis, was not observed. The observed results were consistent with the previous flow cytometry data in Figure 2e which did not show any statistical difference in the percentage of AF488-labelled VLPs taken up by neutrophils characterized by CD11b^+^Ly6G^+^ 3h or 24h post injection in the footpad.

**Table 1.**
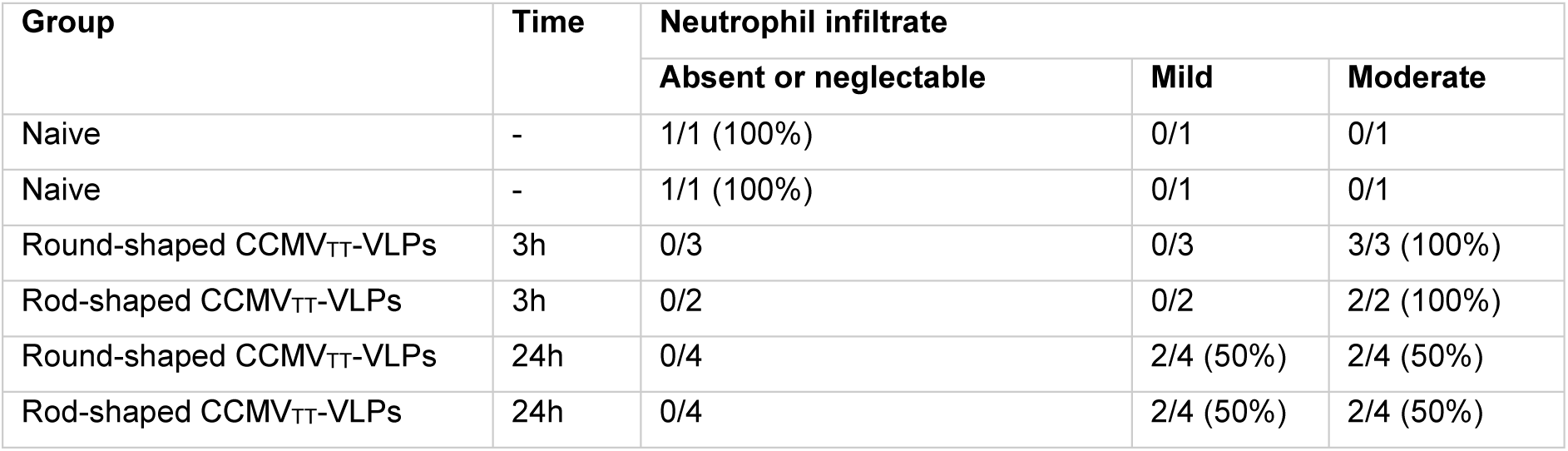
Summary of the relevant histopathological changes in the examined popliteal LNs, indicated per treatment group.

**Figure 7.**
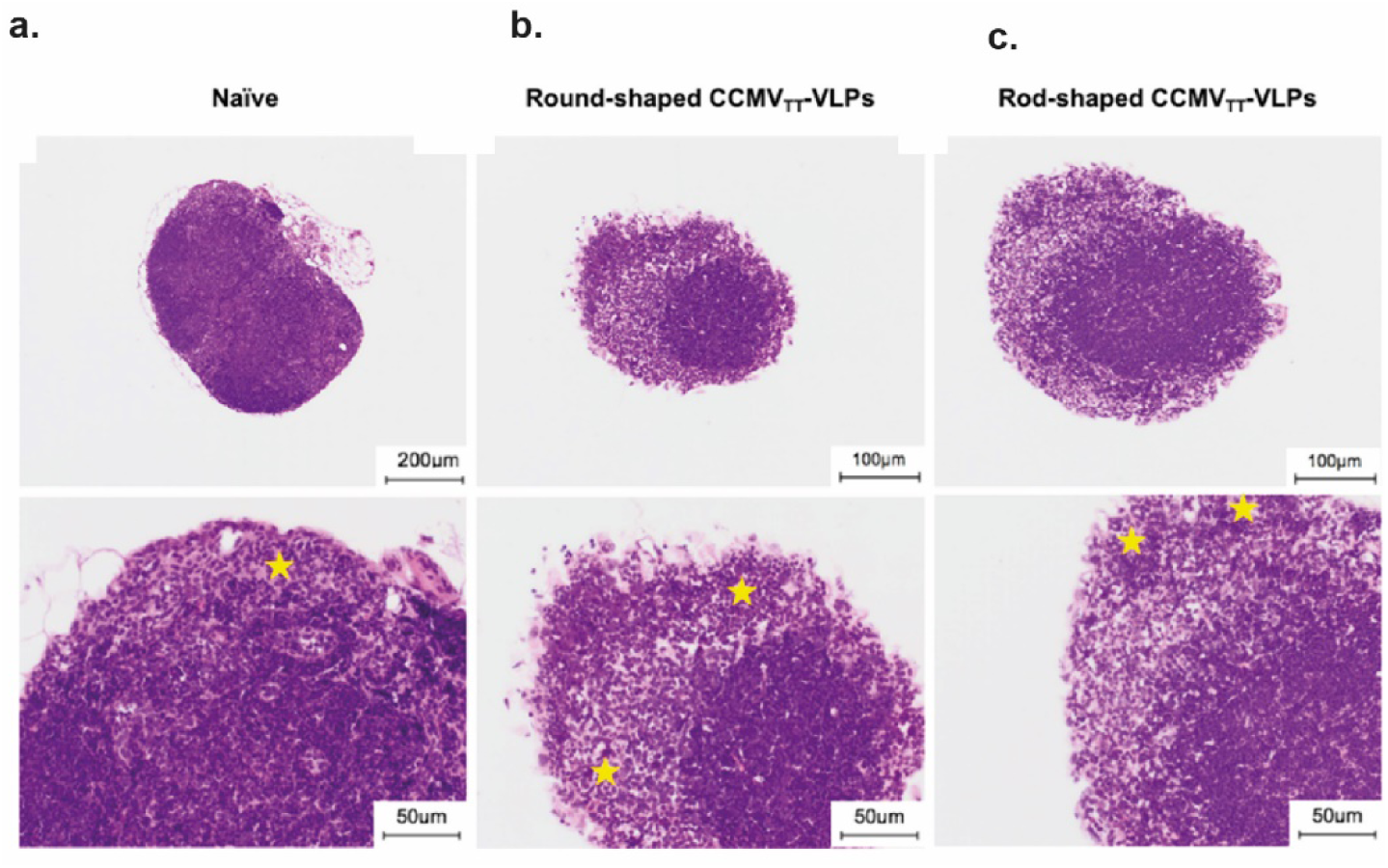
Round and Rod-shaped CCMV_TT_-VLPs enhance neutrophil infiltration in lymph node sinuses without causing any degenerative or vascular changes. ***a***, Top: Naïve popliteal LN 10x objective, HE stain, Bottom: Closer view of Naïve popliteal LN. Within the sinuses (*), granulocytes are rare. 40x objective, HE stain. ***b***, Top: Popliteal LN, 3h after footpad injection with Round-shaped CCMV_TT_-VLPs. 20x objective, HE stain, Bottom: Closer view of popliteal LN, 3h after footpad injection with Round-shaped CCMV_TT_-VLPs. Moderate predominantly neutrophil infiltration in the subcapsular sinus (*). 40x objective, HE stain. ***c***, Top: Popliteal LN, 3h after footpad injection with Rod-shaped CCMV_TT_-VLPs, 20x objective, HE stain. Bottom: Closer view of popliteal LN, 3h after footpad injection with Rod-shaped CCMV_TT_-VLPs. Moderate predominantly neutrophil infiltration in the subcapsular sinus (*). 40x objective, HE stain.

## Discussion

In the current study we have engineered VLPs with nearly identical primary sequence but fundamentally different structural properties; one forming round icosahedrons with a diameter of around 30nm while the other formed rods of about ∼1µm. To this end, we inserted a universal tetanus toxoid (TT) epitope at the N or C-terminus of cowpea chlorotic mottle virus (CCMV)-VLPs. The insertion of TT epitope at the N-terminus did not interfere with the original parental structure and resulted in icosahedral particles T=3; named in this study as Round-shaped CCMV_TT_-VLPs. However, inserting TT epitope at the C-terminus of CCMV_TT_-VLPs caused the formation of Rod-shaped CCMV_TT_-VLPs of ∼1µm in length and ∼30nm in width. As both engineered VLPs were expressed in *E*. coli, they packaged a similar quantity of ssRNA serving as TLR7/8 agonists recognized by PRRs for effective stimulation of the innate immune system. This allowed us to study the impact of size on drainage dynamics and the magnitude of the induced immune responses with one and the same VLP monomer while excluding the effect of TLR7/8 ligands. As mentioned above, both the surface and mass of rods are proportional to the diameter and length, allowing to vary the surface exposed to B cells with a proportional change in mass.

B-cell activation by antigens is a critical step for effective initiation of the adaptive immune response (38). Particulate antigens like VLPs can be passively or actively transported to the lymphoid organs following injection. The passive transportation is based on their ability to drain freely through the lymphatic vessels since they have an ideal size of ∼30-50nm. Our previous studies have proven that icosahedral VLPs such as bacteriophage Qβ-VLPs and VLPs derived from cucumber-mosaic virus (CuMV-VLPs) can freely reach the draining LN in less than a minute, both having a size of ∼30nm (6, 7, 39). By contrast, particulate antigens larger than 500nm cannot efficiently enter the lymphatic capillaries (40, 41). Instead, they need to be actively transported via specialized cells (42). To visualize the trafficking kinetics of the engineered Round and Rod-shaped CCMV_TT_-VLPs (∼30nm and ∼1µm) respectively, they were labelled with AF488 and injected in mouse footpads. Flow cytometric analysis and cryosections of the popliteal LNs 3h and 24h post injection were performed. The results demonstrate more efficient drainage of the Round-shaped CCMV_TT_-VLPs via the lymphatic vessels, which have pores of ∼200nm, 3h following injection in the footpad in comparison with the Rod-shaped ones. Nevertheless, Rod-shaped CCMV_TT_-VLPs have also been detected in the draining popliteal LN 3h post injection in the mouse footpad. This observation may be explained by 2 scenarios: 1) Rod-shaped CCMV_TT_-VLPs exhibit a width of ∼30nm which may allow them to drain into the lymphatic capillaries despite their length. Indeed, if spheres of >500nm size are used for injection, these spheres required 24 hours to arrive in LNs and fully depend on cellular transport (43) and 2) our FACS analysis data indicates that neutrophils characterized by CD11b^+^ Ly6G^+^ can also actively transport both engineered VLPs in a similar manner.

Different subsets of APCs participated in uptake Round or Rod-shaped CCMV_TT_-VLPs upon injection in mouse footpads. Both lymphoid-resident DCs, CD8^+^CD11c^+^ and CD8^-^CD11c^+^, were more efficient at transporting Round-shaped CCMV_TT_-VLPs. Generally, CD8^+^ DCs are more potent in cross-presenting VLP-derived antigens (44). The subcapsular sinus macrophages are considered the frontline cells to capture pathogens in the draining LN and retain them from entering the LN parenchyma. Afterwards they relay the antigen to B-cells for efficient priming and induction of humoral immune responses (45). Yolanda, et al. proposed a model for particulate-antigen acquisition by B-cells. The model suggests that particulate antigens firstly accumulate in the macrophage-niche area in the SCS of the draining LN followed by a still unknown filtration process of the antigens to the follicle. Next, non-antigen specific B-cells carry particulate antigens from the SCS to be deposited on FDCs (39). Our fluorescent microscopy cryosections could demonstrate such findings as Round-shaped CCMV_TT_-VLPs were more efficient in draining to the popliteal LNs 3h post injection in the mouse footpad where macrophages could also be abundantly observed. 24h later the Round-shaped CCMV_TT_-VLPs could be detected deeper in the LN. The binding of the Round-shaped CCMV_TT_-VLPs by B-cells was significantly higher (*p*. <0.0001) when compared to the Rod-shaped CCMV_TT_-VLPs, both at 3h and 24h post injection in the footpad as shown using FACS analysis. This is consistent with our previous observation that Round-shaped VLPs bind to B cells in a complement and CD21-dependent manner (46).

The repetitive surface geometry of VLPs enhance their cross-linking to B-cells and ability to activate complement (38). T=3 VLPs are capable of cross-linking 180 BCRs resulting in a strong humoral immune response. T=3 VLPs may be favorable over T=1 as the later can cross-link ∼60 BCRs which is at the threshold for an optimal immune response (47). However, data are scarce in regard to rod-shaped VLPs and their ability to activate B cells. In this study, we show that a single priming injection of Round-shaped CCMV_TT_-VLPs was efficient at inducing a high specific Ab titer which was further enhanced upon boosting on day 14. On the contrary, the Rod-shaped CCMV_TT_-VLPs could only induce a specific Ab response following boosting on day 14 which remained much reduced compared to the Round-shaped VLP induced response. These results were confirmed by the significantly increased formation of total no. of GCs (*p*. 0.0042) as well as no. of GCs/mm^2^ (*p*. 0.0033) in spleens 12 days following vaccination with Round-shaped CCMV_TT_-VLPs.

When testing the vaccinated mice sera against the opposite engineered VLPs, Round-shaped CCMV_TT_-VLPs were also more efficient in recognizing the Rod-shaped ones even after a single dose. These data indicate that icosahedral VLPs are capable of inducing specific Ab directed against other forms of the same VLPs in an efficient manner and that 30nm sized round VLPs are far superior over 1µm sized rods.

To achieve successful IgG class-switching and memory formation in B cells, co-delivery of innate immune stimuli is crucial (18). It has been shown that class-switching to IgG2a/c and IgG2b is dependent on simultaneous engagement of BCR and TLR9 after immunization with particulate antigens (48, 49). Furthermore, TLR7 engagement with different RNA types influenced the outcome of the humoral immune response to VLP immunization. Bacterial RNA pointed the immune response toward IgG2 production, whereas eukaryotic RNA induced responses favored high IgG1 titers (50). IgG1 is associated with TH_2_ responses, whereas IgG2a/c and IgG3 is associated with TH_1_ responses even though TLR-signaling in B cells is key for IgG subclass induction. The obtained data reveal that Round-shaped CCMV_TT_-VLPs were more efficient than the rod-shaped at inducing class-switching. Furthermore, the ratio between TH_1_ and TH_2_ associated IgG subclasses was more significant (*p*. 0.0045) when vaccinating with Round-shaped CCMV_TT_-VLPs, indicating that rod-shaped VLPs are less effective at driving TLR7-signaling.

Similar to IgG2a responses, VLPs packaged with RNA lead to a strong systemic IgA response. We have shown previously that IgA response is TH cell independent (35) and requires TLR7/8 or 9 to induce a systemic response (37). Here, we show that the systemic IgA response measured on day 21 using a SC prime-boost regimen was much higher in mice vaccinated with Round-shaped CCMV_TT_-VLPs than in mice vaccinated with Rod-shaped CCMV_TT_-VLPs (*p*. <0.0001). Hence, these results also support that TLR7-signaling in B cells is inferior if rods are used for immunization.

The histologic analysis of LNs after immunization with either Round or Rod-shaped CCMV_TT_-VLPs showed an increased number of predominantly neutrophils which were interpreted to correspond to draining leukocytes through the sinuses. The term ‘infiltrate’ is preferred over the term ‘inflammation’ (i.e. lymphadenitis) due to the lack of degenerative or vascular changes in the examined LNs (51, 52). Neutrophil infiltrate was more evident in the tissues sampled 3h than in those sampled 24h post injection, in agreement with the known rapid recruitment of these cells to sites of infection or tissue injury and the short half-lives of neutrophils (53). Increased numbers of neutrophils were absent or neglectable in the examined control LN tissue, which makes a background lesion unlikely. No evident histological difference was observed between the LNs corresponding to the two groups of Round or Rod-shaped CCMV_TT_-VLPs, indicating that both types of VLPs effectively recruit neutrophils.

Taken together, our data demonstrate that antigen size is a key determinant of immunogenicity and that icosahedral antigens of ∼30nm in diameter are far more immunogenic than essentially the same viral capsid protein assembled into µm sized rods. Trafficking from injection site to LNs as well as trafficking within LNs and direct interaction with B cells explain the difference.

## Acknowledgment

This work was supported by Qatar National Research Fund (PDRA grant PDRA4-0118-18002), the Swiss National Science Foundation (SNF 310030_185114 and SNF 310030_179165) and by Latvian Science Council (Grant No. lzp-2019/1-0131).

## Declaration of interests

The authors declare no competing interests

## Notes

### Competing Interest Statement

The authors have declared no competing interest.

