## Supplementary Figures (1 + 2) for "The impact of size on particle drainage dynamics and antibody response"

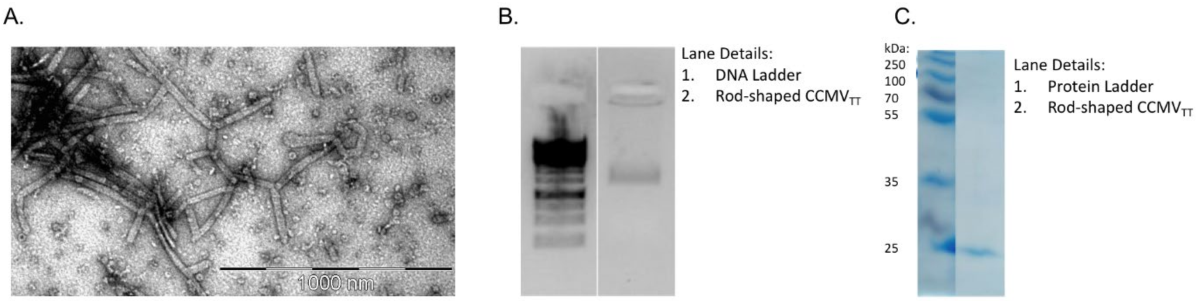


***Supplementary Figure 1: Directional insertion of tetanus toxoid (TT) epitope in the N or C-terminal results in Round or Rod-shaped CCMV$_{TT}$-VLPs. a***, EM of Rod-shaped CCMV$_{TT}$-VLPs with variable fragmented pieces, adsorbed on carbon grids and negatively stained with uranyl acetate solution, scale bars 1000nm. ***b***, Agarose gel stained with SYBR safe, lane 1: DNA ladder, lane 2: Rod-shaped CCMV$_{TT}$. ***c***, Reducing SDS-Page stained with coomassie-blue stain, lane 1: protein marker, lane 2: Rod-shaped CCMV$_{TT}$.

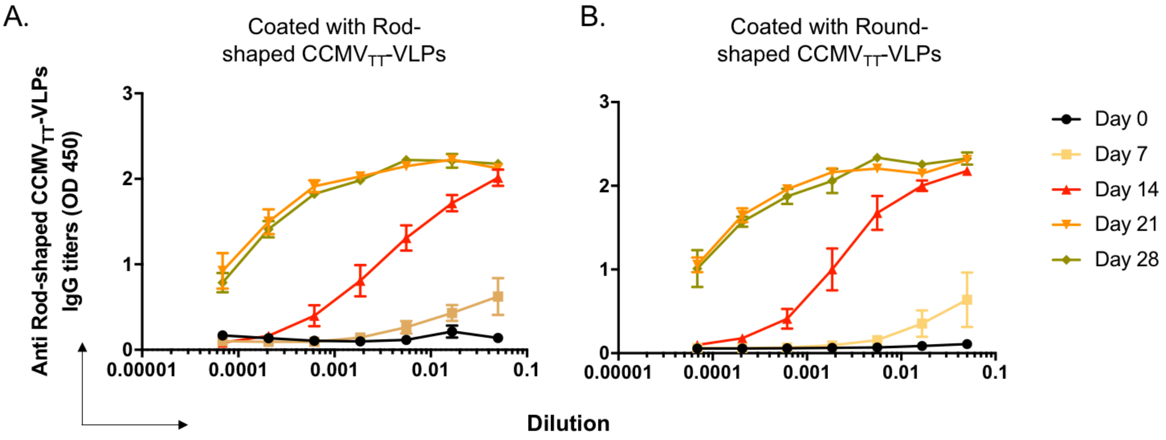


***Supplementary Figure 2: Round-shaped CCMV$_{TT}$-VLPs are more potent in inducing IgG antibodies than Rod-shaped CCMV$_{TT}$-VLPs. a***, Anti Rod-shaped CCMV$_{TT}$-VLPs IgG titers, ELISA plates coated with Rod-shaped CCMV$_{TT}$-VLPs. ***b***, Anti Rod-shaped CCMV$_{TT}$-VLPs IgG titers, ELISA plates coated with Round-shaped CCMV$_{TT}$-VLPs. Mean ± SEM, 3 mice per group, one representative of 2 similar experiments is shown.
